# Rethinking root-shoot growth dynamics

**DOI:** 10.1101/2020.06.29.177824

**Authors:** David Robinson

## Abstract

Using a simple plant growth model based on the logistic equation I re-evaluate how biomass allocation between roots and shoots articulates dynamically with the rate of whole-plant biomass production. Defined by parameters reflecting lumped physiological properties, the model constrains roots and shoots to grow sigmoidally over time. From those temporal patterns detailed trajectories of allocation and growth rate are reconstructed. Sigmoid growth trajectories of roots and shoots are incompatible with the dominant ‘functional equilibrium’ model of adaptive allocation and growth often used to explain plants’ responses to nutrient shortage and defoliation. Anything that changes the differential rates of growth between roots and shoots will automatically change allocation and, unavoidably, change whole-plant growth rate. Biomass allocation and whole-plant growth rate are not independent traits. Allocation and growth rate have no unique relationship to one another but can vary across a wide spectrum of possible relationships. When root-shoot allocation seems to respond to the environment it is likely to be a secondary illusory consequence of other primary responses such as localised root proliferation in soil or leaf expansion within canopy gaps. Changes in root-shoot allocation cannot themselves compensate directly for an impairment of growth rate caused by an external factor such as nutrient shortage or defoliation; therefore, such changes cannot be ‘adaptive’.

‘The reasons are so simple they often escape notice.’ (James 2012, p. 6).

## Introduction: Handcuffing an octopus

A doyen of plant growth analysis once told me that trying to understand how a plant’s biomass is allocated between roots and shoots while it simultaneously produces more biomass is like ‘trying to handcuff an octopus’: no sooner do you pin down whole-plant growth rate than allocation slips out of your grasp; nail allocation and you find growth rate has hidden itself under a rock. At first sight it may seem unlikely that this is so, given that the production of biomass and its allocation among different organs are such well-studied phenomena. Nevertheless, some surprisingly simple aspects of how these processes are related have been overlooked or misunderstood.

Thinking on this subject is dominated by Brouwer’s ‘functional equilibrium’ concept of plant growth (Brouwer 1962). Admirably reviewed by Poorter & Nagel (2000), Brouwer’s theory was encapsulated thus: ‘… plants shift their allocation towards shoots if the carbon gain of the shoot is impaired by a low level of above-ground resources, such as light and CO_2_ [and also if the plant is defoliated: DR]. Similarly, plants shift allocation towards roots at a low level of below-ground resources, such as nutrients and water. These shifts could be seen as adaptive, as they enable the plant to capture more of those resources that most strongly limit plant growth’ and later: ‘…implicit in this model is that a plant allocates its biomass in such a manner that its growth rate is maximal under the given environmental conditions.’ This intuitively attractive idea underpins many plant growth models (see Wilson 1988; Thornley 1998).

The key assumption is that there are optimum allocations of biomass at which the rate of biomass production is maximised. But if the key assumption is true, why has it apparently been impossible for anyone to plot measurements of whole-plant growth rate against some measure of allocation to see the former maximised at a certain value (or values) of the latter? Why is the relationship between allocation and growth rate so enduringly enigmatic?

Part of the problem is that few relevant experiments have followed root and shoot growth for long enough and in enough detail to reveal the full dynamics of biomass allocation and its co-variation with biomass production rates. Complex growth models have not revealed, or been used to reveal, just how biomass allocation articulates dynamically with growth rate. Maybe it seems too trivial an issue to attract the firepower of a full-blown simulation model, or the models are just too specific to clarify what is, in my view, a very general and important relationship.

Whatever the reasons, we need an alternative approach to resolve how root-shoot allocation and whole-plant growth rate co-vary. That which I propose here takes its cue not from plant physiology, but from population biology, a field dominated by simple phenomenological models rather than mechanistically realistic ones.

### Simplifying the complex

#### Preamble

The starting point for this re-think is the seemingly trivial observation that root and shoot growth change with time in both absolute terms and relative to one another, and that those changes can to a reasonable approximation be described mathematically without worrying too much about the fine details.

Trajectories of root and shoot growth can approximate simple linear or exponential functions of time in young plants (the subjects of almost all the influential experiments on root-shoot allocation) or they can be described with ever increasing statistical precision by whatever polynomial or other curve best fits the data. But the most general temporal pattern of root and shoot growth is a sigmoid trajectory.

All plants grow sigmoidally over much of their lives even if, in the short-term and when young, they might seem to grow linearly or exponentially. If they do, then they cannot do so forever. Models using equations describing mechanisms of light interception, nutrient uptake, organ formation and resource allocation (for example, Brown *et al.* 2019) always predict approximately sigmoid trajectories of biomass change. Biomass production eventually reaches a ceiling of one sort or another. That ceiling may be only a transient halt before regrowth or reproduction occurs, or it can be a prelude to senescence and death. The timespan of a sigmoid trajectory will range from days or weeks for a fast-growing short-lived annual species, to decades or centuries for a slow-growing long-lived tree. A sigmoid growth pattern is as close to a universal biological principle as any (West *et al.* 2001).

If separate trajectories of roots and shoots are constructed, they can be combined to calculate biomass allocation between roots and shoots along with the growth rate of the whole plant (Robinson & Peterkin 2019). This allows the co-variation between biomass allocation and production to be visualised directly in ways that existing models or experiments have not. If the supposed ‘adaptive’ responses of allocation referred to above are real, then they should appear in such visualisations.

There are strong arguments that the root-shoot dichotomy is just too simple to capture the reality of plant growth. At least three classes of structures – leaves, stems and roots – are needed to reflect their roles in above- and below-ground resource capture as well as mechanical support of a vegetative plant (Poorter & Nagel 2000; Poorter *et al.* 2015). Biomass allocation in a reproductive angiosperm would also need to account for flowering stalks, sexual organs, flowers, fruit and seed. And distinguishing coarse roots from fine roots reflects more faithfully what those structures contribute to the life of their parent plant (Chen *et al.* 2019). While recognising that it caricatures the behaviour of real plants, I have retained the classical root-shoot division because the loss of realism is offset by gains in clarity and generality.

#### Producing biomass

Incremental changes in root or shoot biomass between two successive times, *t*_1_ and *t*_2_, and which form part of a sigmoid trajectory can be described by an equation such as the logistic. A mainstay of population modelling (‘probably the simplest nonlinear equation one could write’: May 1974a), the logistic equation can be written as:

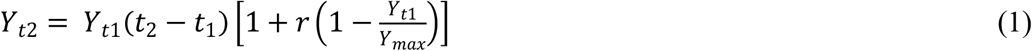

where *Y* is root (*R*) or shoot (*S*) biomass (in terms of dry weight) of an individual plant, *Y_max_* the asymptotic maximum value of *Y, t* time, and *r* the intrinsic growth rate of *Y* (with units of [time]^-1^). If the time interval *t*_2_ – *t*_1_ is sufficiently short and many such successive intervals are assumed, the difference equation, eqn. (1), approximates a continuous estimate of *Y* over time. There is nothing special about eqn. (1). Any function that produces a sigmoid trajectory of *Y* with time could be used for this purpose although the algebraic details will differ.

For simplicity, I assume that *Y_max_* and *r* are independent of one another, and root-specific values of *Y_max_* and *r* independent of those for shoots. In real plants, root and shoot growth and other physiological processes are interdependent to some extent, and the activities of one plant part feeds back to influence the activities of another. Roots and shoots are connected by common vascular systems and the continual exchange between them of numerous resources and signals help coordinate their respective activities. Experiments on young seedlings often give the impression that roots and shoots do grow synchronously. It is a mistake, however, to suppose this the norm. One look at the disparity between the above- and belowground phenologies of numerous species dispels the notion of there always being close synchrony between root and shoot growth (Abramoff & Finzi 2015; Makoto *et al.* 2020).

Whole-plant biomass (*W*) at any time is simply the sum of *R* and *S* at that time, and its value depends on how eqn. (1) is parameterised separately for roots and shoots in terms of *r* and *Y_max_* values. In addition, when using eqn. (1) in the absence of experimental data to which the curve can be fitted (for example, Robinson & Peterkin 2019), it is necessary to specify initial values of root and shoot mass (*R*_0_ and *S*_0_). It is more likely than not that initial root and shoot masses will differ, so that is what I assume here.

The absolute growth rate (AGR, with units of [mass] [time]^-1^) of root or shoot biomass at any point is the first derivative of eqn. (1) with respect to time, so that AGR = *rY* (1 - *Y* / *Y_max_*) where, again, *r*, *Y* and *Y_max_* are specific for roots and shoots.

This definition of AGR says that the instantaneous growth rates of roots and shoots depend on the values of both *r* and *Y_max_*. The influence of *r* on growth rate is greatest when *Y* is small compared with *Y_max_*, that is, when *Y* is much smaller than *Y_max_* AGR approximates *rY*. Conversely, when *Y* is close to its upper limit *r* can have little influence on growth rate because then AGR is virtually zero whatever the value of *r*. In other words, the value of *r* has most influence on young plants and none on old plants. *Y_max_* has negligible influence on young plants. Its main influence is to determine how long it takes *Y* to reach its ultimate value: the bigger *Y_max_*, the longer it takes. *r* and *Y_max_* exert their most powerful joint influences on root or shoot growth rate somewhere in the middle of their sigmoid trajectories.

The growth rate of a whole plant or of its component parts is usually expressed as its relative (or specific) growth rate (RGR, with units of [time]^-1^). Once values of *R* and *S* have been calculated by eqn. (1) it is straightforward to calculate RGR separately for roots, shoots and for the whole plant between successive time intervals *t*_1_ and *t*_2_ (Hunt 1982, p. 18).

Whole-plant RGR is:

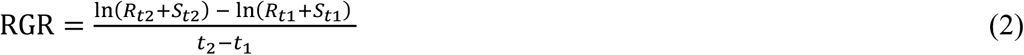

where, as before, *R* and *S* are root and shoot biomasses, respectively. Root and shoot RGRs are calculated in the same way, leaving out the shoot or root biomasses from eqn. (2), as appropriate. Again, if *t*_2_ – *t*_1_ is short and there are many such intervals eqn. (2) approximates a continuous estimate of RGR with respect to time.

RGR of the whole plant is also defined instantaneously as AGR/*W*. From the above definition of AGR, RGR = *r* (1 - *Y* / *Y_max_*), where *Y* = *W*=*R* + *S* as calculated by eqn. (1), *Y_max_* = *W_max_* = *R_max_* + *S_max_*, and *r* in this case is the weighted mean of the *r* values for roots and shoots. Therefore, RGR equals *r* at zero biomass (*Y* = 0) or, equivalently, at zero time. The maximum possible RGR is *r*. RGRs of roots and shoots inevitably decline over time as *Y* gets closer to *Y_max_*, unless that progression is interrupted by, for example, sudden biomass loss in a defoliation event. Because whole-plant RGR is calculated from the combined trajectories of *R* and *S,* it is not necessarily true that whole-plant RGR will always decline with time. The temporal trajectory of whole-plant RGR will depend on the values of *r* and *Y_max_* assumed for roots and shoots.

#### Allocating biomass

Root-shoot biomass allocation at any time is expressed most simply as the plant’s root mass fraction (RMF):

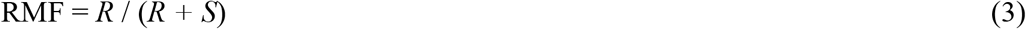

where *R* and *S* are each calculated separately by eqn. (1). Because, via eqn. (1), both *R* and *S* are functions of time, so is RMF. RMF can have any value between 0 and 1, but physiological constraints limit it to somewhere within that range. RMF = 0.5 indicates equal biomass allocation to roots and shoot; greater than 0.5 means more biomass is allocated to roots, and less than 0.5, the opposite. In most plants RMF is typically between 0.1-0.4 with a global median of 0.24 (based on 11217 data from 1208 species in Table S2 of Poorter *et al.* 2015), although the extremities of that database range from an RMF as small as 0.02 in *Manilkara bidentata* to one as large as 0.92 in *Picea abies.*

For a plant whose growth trajectory follows eqn. (1) to ‘adaptively’ change its root-shoot allocation of biomass it must change the RGRs of its roots, shoots or both (Brouwer 1962). Because RGR = *r* (1 - *Y* / *Y_max_*), that means changing *r*, *Y_max_* or both. If root and shoot RGRs cannot change, allocation cannot deviate from its current trajectory. Of course, ‘changing *r* and *Y_max_*’ is just shorthand for the myriad molecular and genetic processes that regulate biomass production, notable recent examples of which include the nitrogen-induced increase in abundance of growth-regulating transcription factors (Swift *et al.* 2020) and the temporal dynamics of multiple QTL associated with RGR (Meyer *et al.* 2020). But for this exercise, such processes and their mechanisms need not be specified – fortunately for us.

Another simplification I make is to ignore the many complications thrown up by clonal or parasitic plants, plants that go into long periods of dormancy, interannual variations in the growth of long-lived plants, the influences of symbionts, and competing plants, to list just a few of the reasons why what follows aims ‘not at realism in detail, but rather at providing mathematical metaphors for broad classes of phenomena. Such models can be useful in suggesting interesting experiments or data collecting enterprises, or just in sharpening discussion’ (May 1974b, p. *v*) – *especially* in sharpening discussion.

### Starting with some simple sigmoid trajectories

Let’s start with a simple example to illustrate some basic points. I assume here that our plant grows with certain arbitrary values of *r* and *Y_max_* for its roots and shoots: for roots *r* = 0.06 and *Y_max_* = 30; for shoots *r* = 0.1 and *Y_max_* = 50. Assume, again arbitrarily, that the initial root mass (*R*_0_) = 0.8 and the initial shoot mass (*S*_0_) = 1.2, so that the initial total plant mass = 2. The resulting growth trajectories are plotted in Fig. 1 over 100 time-steps.

**Fig. 1.**
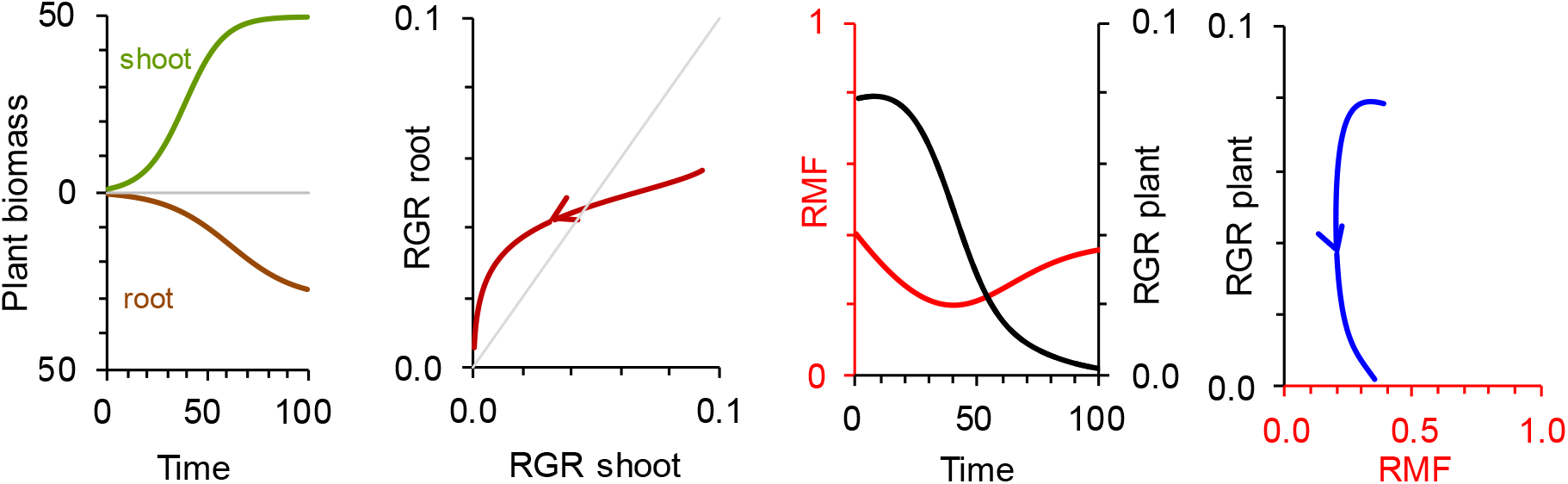
Trajectories of root and shoot biomass with time (eqn. (1)), root, shoot and whole-plant RGRs (eqn. (2)), and RMF (eqn. (3)). The curves in the third panel are plotted against each other in the fourth. Throughout, root *r* = 0.06 and *Y_max_* = 30, shoot *r* = 0.1 and *Y_max_* = 50. Initial root mass (*R*_0_) = 0.8 and initial shoot mass (*S*_0_) = 1.2. Arrows indicate temporal direction.

The sigmoid progressions of root and shoot masses generate continuously declining RGRs of roots and shoot. Shoot RGR initially exceeds that of roots, but these later reverse and each eventually declines towards zero. Because in this example root and shoot RGRs are always changing it is inevitable that root-shoot biomass allocation, as embodied in the root mass fraction, also changes continually. This change in RMF occurs in the absence of any external driver which makes RMF rise or fall.

The temporal trajectory of RMF in Fig. 1 reflects its ‘ontogenetic drift’ (Wilson 1988) or ‘apparent plasticity’ (McConnaughay & Coleman 1999). Ontogenetic drift in biomass allocation arising from a disparity in growth rate between roots and shoots is possible even if the parameters defining root and shoot growth rates, namely *r* and *Y_max_*, are constant. A phenotype fixed in terms of how eqn. (1) is parameterised can still exhibit plasticity. The prospect of a plant adjusting its growth to maintain something like a steady root-shoot biomass allocation over time seems improbable if RMF is such an intrinsically labile quantity. It is true, however, that small temporal changes in RMF might be barely detectable in practice against the background variation among experimental replicates.

Towards the end of the time-course when neither root nor shoot biomass changes much (because each is close to its *Y_max_*) RMF approaches a steady value. Only if root and shoot RGRs are equal and constant will allocation stay constant over time. That will be a rare occurrence.

The corresponding whole-plant RGR in this example does not decline continually, unlike the RGRs of its component root and shoot masses. It initially increases slightly, reaching its maximum at *t* = 7, before then declining towards zero as roots and shoot approach their upper limits.

Plotting whole-plant RGR against RMF produces a co-trajectory that I doubt many of us would have predicted from existing theoretical knowledge or empirical evidence. As far as I know such a trajectory has yet to be discovered by any experiment, although by plotting partial RGR-RMF trajectories derived by fitting eqn. (1) to biomass measurements of 11 herbaceous species Robinson & Peterkin (2019) hinted at what Fig. 1 shows in full.

Perhaps the most important feature of the RGR-RMF relationship in Fig. 1 is that although in this case there does happen to be an optimum RMF (RMF = 0.34) associated with a maximal RGR (RGR = 0.0793) at *t* = 7, that is simply a transient point generated by the particular values assumed for the variables. There is no possibility of the plant somehow adjusting its RMF to maximise its RGR by returning to those coordinates while preserving the sigmoid trajectories of roots and shoots; it is not an equilibrium towards which a growth trajectory could converge in any apparently goal-seeking way (Thornley 1998).

In the long run the RGR of an individual must decline, although, as Fig. 1 suggests and experimental evidence (Hunt 1982, pp. 20, 192; Hunt & Lloyd 1987) proves, RGR can initially increase from zero (or even from negative values) to a maximum before falling gradually. The cause of this pattern is probably nothing more than a difference in growth rate between roots and shoots during post-germination development.

The full richness of the how RGR and RMF co-vary can be explored by plugging into eqn. (1) different values of *r* and *Y_max_*, but I will defer that until later, after first considering what happens to a plant when part of it gets eaten.

### Taking a big bite of biomass

If the plant whose growth is shown in Fig. 1 is defoliated by allowing a well-trained rabbit to instantly eat nine-tenths of its shoot biomass at *t* = 50, the result is as shown in Fig. 2. Until *t* = 50, of course, everything proceeds as before. Afterwards, nothing is the same.

**Fig. 2.**
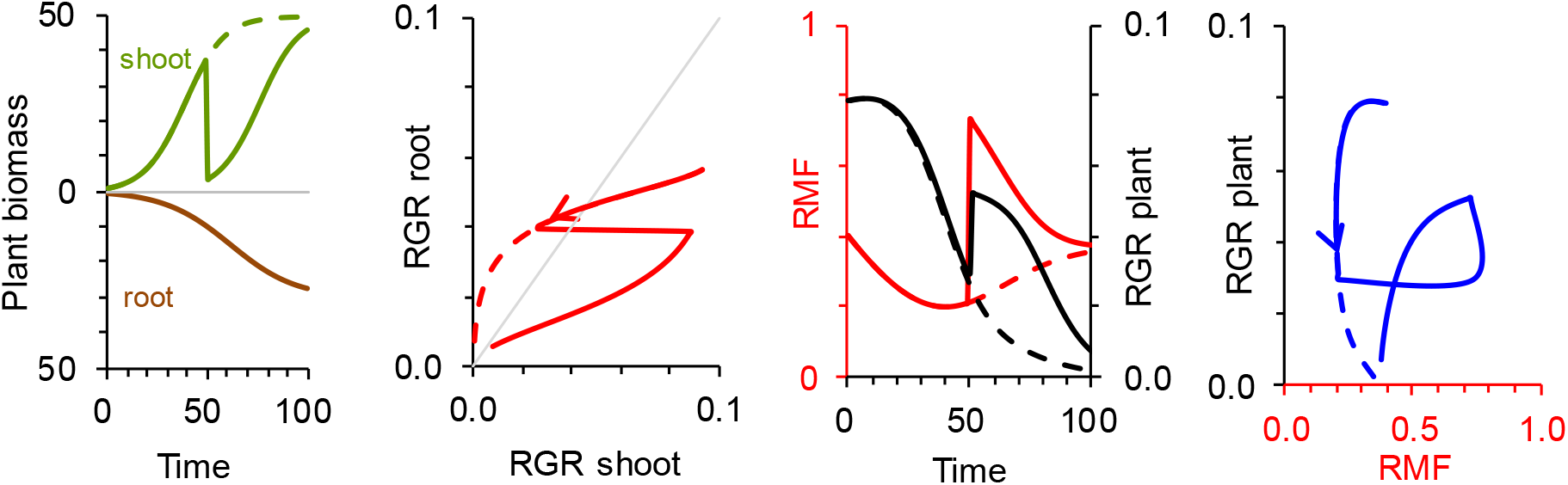
As for Fig. 1 but at *t* = 50, 90% of shoot biomass is removed in a defoliation event. *r* and *Y_max_* values are the same throughout, as in Fig. 1. For comparison, the trajectories of the undefoliated plant in Fig. 1 are included here as broken lines.

Obviously, when defoliated the plant’s RMF shoots upwards from its previously smooth progression. Eventually, RMF declines back towards where it would have been had the plant remained intact. Interestingly, so does RGR. For the remainder of the time-course, the RGR of the defoliated plant comfortably exceeds that of its unmolested counterpart. The latter plant is, of course, bigger than the former, but it grows more slowly. So, there is a possibility here for an increase in RMF following defoliation being associated with a temporarily faster RGR compared with an undefoliated plant. That effect has been seen in some clipping experiments (for example, van Staalduinen & Anten 2005) but it is not universal, any effect in real plants depending strongly on context, that is, species, clipping regime and environmental conditions (Hilbert *et al.* 1981; Coughenour 1991).

Conventional physiological wisdom might interpret the post-defoliation increase in RMF in Fig. 2 as an ‘adaptive response’ giving a compensatory boost to whole-plant RGR, and the restoration of an equilibrium implied by the eventual return of root and shoot trajectories close to where they would have reached anyway. But that interpretation would be wrong.

The faster post-defoliation RGR in Fig. 2 is a direct, if temporary, artefact of whole-plant biomass being smaller after defoliation than before. Because RGR = *r* (1 - *Y* / *Y_max_*), a smaller *Y* (whole-plant biomass in this case) produces a faster RGR after *t* = 50 even if *r* and *Y_max_* remain constant. Both *r* and *Y_max_* are the same before defoliation and after. In that sense, the plant’s growth cannot be said to have ‘responded’ physiologically to part of it being eaten. No physiological mechanisms of compensation are needed to explain the plant’s growth dynamics after defoliation.

No such mechanism could explain in any case the RGR-RMF co-trajectory shown in Fig. 2. As bizarre it seems, that trajectory originates directly from the continued unfolding of sigmoid growth of roots and shoots even when determined by fixed parameter values, and not from any special response to defoliation and certainly not a response that converges towards anything identifiable as an optimum.

However, a real plant well might alter its *r* or *Y_max_* values by triggering specific physiological mechanisms in response to rabbit attack. For example, defoliation can slow root growth (Wilson 1988), an effect not included in the simulation in Fig. 2 where constant root *r* and *Y_max_* are assumed. Slower root growth in a defoliated plant implies, in terms of the sigmoid growth model, a reduction in root *r* or *Y_max_* specifically in response to defoliation. If that happened the effects on whole-plant RGR illustrated in Fig. 2 could be amplified or dampened. Fig. 2 shows that specific response mechanisms are not necessary to accelerate growth post-defoliation, but such responses might occur. How could you tell if they did?

One way would be to compare eqn. (1) (or whatever is your preferred model) to plant growth data with and without defoliation, to test if it is necessary to adjust the equation’s parameter values – that is, of *r* and *Y_max_* – to get the best statistical fit. If adjustments in parameter values are necessary, that would be evidence for a genuine physiological response; if not, not.

### Responding to reduced nitrogen (or phosphorus, potassium, sulphur…) availability

When an essential nutrient such as nitrogen suffers a reduced availability, the classical theory says that root growth increases and shoot growth decreases in response. If we simulate this, what happens?

Taking the plant in Fig. 1, at *t* = 25 flip its root and shoot *r* and *Y_max_* values as if its nitrogen supply was decreased at that time. After *t* = 25, root *r* = 0.1 and *Y_max_* = 50, and shoot *r* = 0.06 and *Y_max_* = 30, for the purposes of illustration. The result is in Fig. 3 with, again, the trajectories shown originally in Fig. 1 included for comparison.

**Fig. 3.**
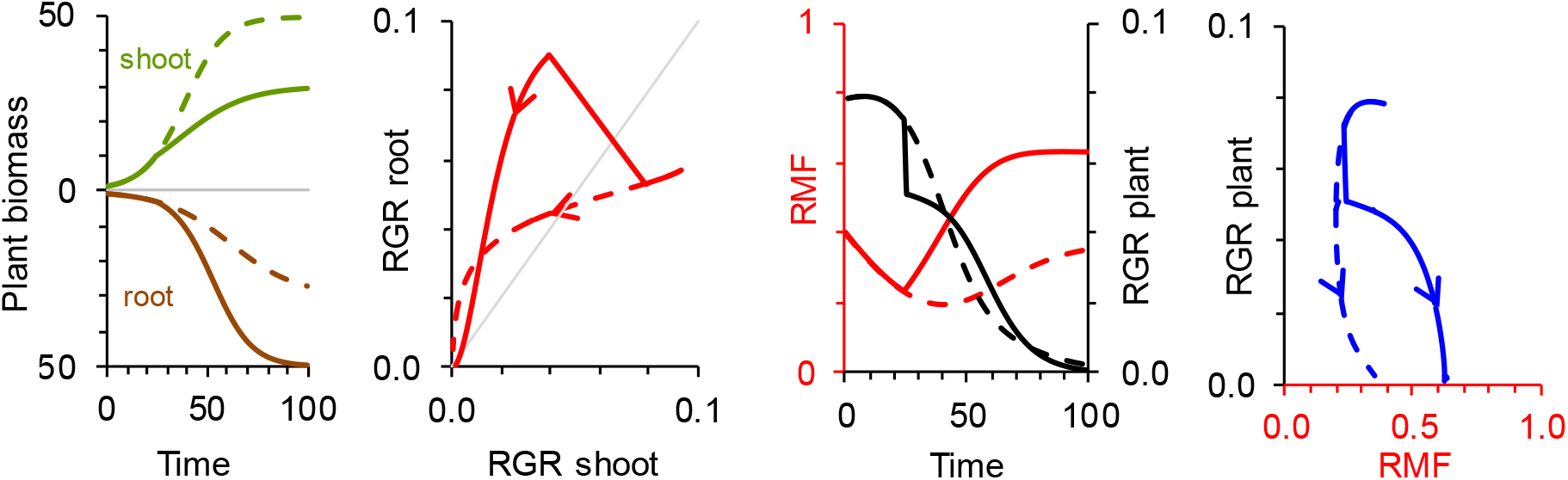
As for Fig. 1 but after *t* = 25, root *r* = 0.1 and *Y_max_* = 50, and shoot *r* = 0.06 and *Y_max_* = 30 to simulate the possible effects of a reduction at *t* = 25 in the availability of an essential nutrient. For comparison, the trajectories of the plant in Fig. 1 are shown here as broken lines.

By changing the values of root and shoot *r* and *Y_max_* in this way, the balance of biomass production after *t* = 25 shifts from shoot to root, as the classical theory says it should. In a real plant the larger root system would then be available to mop up the scarce soil nitrogen to satisfy the relatively smaller metabolic demands imposed by the shoot. This is the physiological rationale for of the potential compensatory effect of such a biomass allocation response to scarcities of nitrogen and other nutrients.

Even though the changes in *r* and *Y_max_* are instantaneous when in a real plant they would probably have occurred more gradually, the resulting change in RMF after *t* = 25 proceeds smoothly, progressively increasing biomass allocation to the roots. What does this do to RGR?

At *t* = 25 there is the immediate reduction in RGR that nitrogen deprivation would likely cause in a real plant. But then RGR rebounds later to exceed, albeit only slightly in this example, the RGR of the phenotype with fixed *r* and *Y_max_* values. Does the dramatic change in RMF after *t* = 25 allow RGR to eventually maintain a faster rate than it could otherwise have done, as expected from the functional equilibrium concept? No, I don’t think it does, for the following reason.

The rebound in RGR is explained again by that plant’s biomass being smaller than it would have been without having its root and shoot *r* and *Y_max_* values switched. The faster RGR in the nitrogen-deprived plant after about *t* = 40 is an artefact of how that quantity is calculated (eqn. (2)). But what is revealing is that similar whole-plant RGRs can be associated with such very different RMFs. This suggests that RGR is only weakly dependent on the value of RMF, contrary to the notion that an optimum biomass allocation is decisive in determining the whole-plant’s growth rate. Why might that be?

To answer this question, we need to explore the wider landscape within which RGR and RMF can co-vary.

### Filling phenotypic space with plastic plants

How much of the RGR-RMF ‘phenotypic space’ (Pigliucci 2007), ‘phenotypic landscape’ (Williams *et al.* 2013) or ‘viable trait space’ (McCormack & Iversen 2019) can be occupied? Do RGR and RMF co-vary in similar ways across that space? And are certain combinations of RGR and RMF off-limits for a plant whose growth follows a sigmoid trajectory, or are virtually any combinations of RGR and RMF feasible?

To tackle these questions, I varied root and shoot *r* and *Y_max_* values as described in the legend to Fig. 4, keeping them constant throughout, so assuming no malign influences of herbivore or environment during growth. In other words, by assuming a wider range of *r* and *Y_max_* values the RGR-RMF co-trajectory shown already in Fig. 1 is joined by others to see where and how they occur. The left-hand panel of Fig. 4 shows the result.

**Fig. 4.**
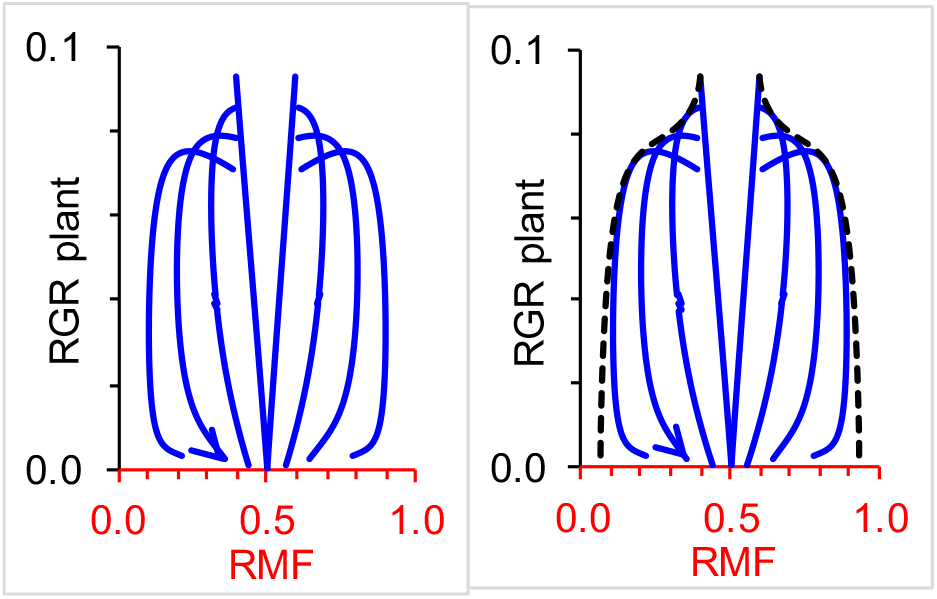
Co-trajectories of whole-plant RGR and RMF generated by varying values of *r* and *Y_max_* that remain constant over time. From left to right in the first panel: root *r* = 0.04, *Y_max_* = 20, shoot *r* = 0.1, *Y_max_* = 50; root *r* = 0.06, *Y_max_* = 30, shoot *r* = 0.1, *Y_max_* = 50 (this, arrowed, is the RGR-RMF co-trajectory shown previously in Fig. 1); root *r* = 0.08, *Y_max_* = 40, shoot *r* = 0.1, *Y_max_* = 50; root *r* = 0.1, *Y_max_* = 50, shoot *r* = 0.1, *Y_max_* = 50 (this is the fourth plot from the left, the linear trajectory generated because *r* and *Y_max_* values are the same for roots and shoots, which is why it converges eventually on RMF = 0.5). Initial root and shoot masses are *R*_0_ = 0.8 and *S*_0_ = 1.2 for all four trajectories. The other four trajectories in the left-hand panel are for plants with initial root and shoot masses reversed, *R*_0_ = 1.2 and *S*_0_ = 0.8, and with *r* and *Y_max_* values that mirror those of the first four, namely, from left to right: root *r* = 0.1, *Y_max_* = 50, shoot *r* = 0.1, *Y_max_* = 50; root *r* = 0.1, *Y_max_* = 50, shoot *r* = 0.08, *Y_max_* = 40; root *r* = 0.1, *Y_max_* = 50, shoot *r* = 0.06, *Y_max_* = 30; root *r* = 0.1, *Y_max_* = 50, shoot *r* = 0.04, *Y_max_* = 20.

The second panel shows the same eight RGR-RMF co-trajectories as those in the first but superimposed on them are two other curves (broken lines) that approximately maximise RGR for a given RMF. The left-hand broken curve was generated by fixing shoot *r* = 0.1 and *Y_max_* = 200 throughout, while making root *Y_max_* = 50 but root *r* vary at each time step according to 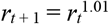. The right-hand broken curve mirrors the first one since it was generated by fixing root *r* = 0.1 and *Y_max_* = 200 throughout, while making shoot *Y_max_* = 50 but shoot *r* vary according to 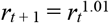.

I was surprised to see so much of the RGR-RMF space in Fig. 4 occupied (and, with a pleasing resonance, occupied in a vaguely octopodal way). It seems that only fast RGRs combined with extreme RMFs are out of bounds. Depending on how the equations are parameterised, RGR can vary positively, negatively or hardly at all with RMF.So, it is unsurprising that the relationship between a plant’s production and allocation of biomass has remained so enigmatic, and that experiments yield results difficult to reconcile with a functional equilibrium. Fig. 4 shows that the same whole-plant RGRs occur in plants with very different biomass allocations. There is no evidence that certain ranges in RMF are intrinsically more conducive to promoting faster RGRs than others. If that’s true, there can be no equilibrium or optimum allocations towards which a plant can grow to maximise its overall growth rate while maintaining a sigmoid growth pattern whatever its *r* and *Y_max_* values might be. This questions the key assumption of the functional equilibrium concept.

It is interesting that RGR-RMF trajectories in Fig. 4 cross-over one another in the early stages of growth. This means that some phenotypes, as defined by fixed *r* and *Y_max_* values, could temporarily out-grow others. But could another phenotype outgrow all the fixed phenotypes by, for example, varying its *r* or *Y_max_* values continually as it grows? At least one phenotype could.

I found this plastic phenotype by trial-and-error. By making shoot *Y_max_* a fixed value of 200 and reducing root *r* at each time step from an initial value of 0.1 according to a power function 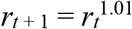 or, by making root *Y_max_* = 200 and reducing shoot *r* at each time step using the same function, the resulting phenotype (the broken curves in Fig. 4) outgrows any of the fixed phenotypes for any RMF attained by them. The RGR of the plastic phenotype still declines over time but not as rapidly as in the other phenotypes. (The idea of changing *r* continually to simulate plant growth via logistic models was used previously by Wallach & Gutman (1976). They modelled, with mixed success, biomass production by communities of winter annuals in arid environments by varying *r* not as a direct function of time, but as arbitrary functions of soil moisture, radiation and temperature.)

I use the arbitrary function 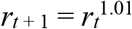 only to demonstrate that at least one continually plastic phenotype is possible within this framework that is able to afford some temporary advantage – in terms of attaining a faster RGR for a given RMF – compared with a fixed phenotype; no doubt there are other plastic phenotypes described by other functions, but I have not explored what they might be.

Whether that plastic phenotype is physiologically possible is another matter. The extent to which I had to contrive it into existence on my spreadsheet suggests it’s unlikely to be realistic. But such an improbably plastic phenotype is what would be needed for genuinely ‘adaptive’ allocation responses to exist in the landscape of opportunity defined by eqns. (1)(3). Even so, the possibility of a phenotype capable of continually changing its root and shoot growth trajectories should not be dismissed entirely when analysing experimental data.

## Discussion

If the trajectories presented here are valid mathematical metaphors for real plants, adjustments in root-shoot allocation cannot themselves compensate whole-plant growth rate for nutrient shortages or defoliation. The possible co-variations between allocation and growth rate are incompatible with the classical functional equilibrium model.

A mistake many of us have made is to think of root-shoot biomass allocation and biomass production rate as if they are independent traits. We have tried and largely failed to understand them in those terms. But they are not independent traits. Just as you can’t change the area of a circle without automatically changing its diameter, biomass allocation can’t change without also changing biomass production rate to a greater or lesser extent. This fundamental misunderstanding is why the metaphorical octopus has taken so long to handcuff. In some ways this parallels the better-known error of attributing allometric changes in root and shoot size as genuine allocation responses to the environment (Reich 2002).

In plant physiology we are used to thinking of our measured variables changing in approximately deterministic ways regulated by well-characterised mechanisms. We understand why the rate of CO_2_ fixation varies with irradiance as it does because we can interpret data with linear models of the essential biochemical and biophysical mechanisms. We can write a simple linear equation describing concentration-dependent ion uptake rate by a root. But we can’t write a simple linear equation describing how a plant’s biomass production rate depends on how it allocates biomass between roots and shoot because there is no independent variable in that relationship. The wide spectrum of possible variation between RGR and RMF in Fig. 4 illustrates this co-dependence, presuming the trajectories in Fig. 4 to be valid, of course. And the way to test their validity is to do appropriate experiments and apply eqns. (1)-(3) to the data. This isn’t the only example in plant physiology of two important variables having no unique relationship with one another. A comparable co-dependency exists between simultaneous rates of CO_2_ assimilation and water evaporation via stomata (Cowan & Farquhar 1977).

So, how does plant growth respond to the environment if not primarily by alterations in root-shoot allocation?

A plant’s environment is never uniform in space or time. Different sets of meristems of the same plant can experience different external cues and conditions. Some growth responses to such an environment do have the potential to be genuinely ‘adaptive’ in terms of compensating for resources distributed non-uniformly and perhaps unpredictably in space and time, or for the partial destruction of biomass. These include localised root proliferation in transiently nutrient-rich soil (Robinson 1994) and leaf and stem expansion into better illuminated parts of the canopy (Küppers 1994). As they occur and as they help the plant to capture scarce resources such responses will almost certainly change differential root or shoot growth rates. If that happens, simultaneous alterations in root-shoot biomass allocation and whole-plant growth rate will be inevitable if the growth of a real plant is caricatured faithfully by eqn. (1).

The changes in root-shoot allocation will appear as if *they* are the primary responses to the environment. They, however, are likely to be only secondary consequences of localised, temporary, and to some extent independent growth responses of roots and shoots to nutrients, light, herbivory and so on. It is *those* potentially ‘adaptive’ responses that cause root and shoot growth rates to differ and whole-plant growth rate to change; that’s the illusion that has been fooling us for so long.

But such changes will occur whenever anything – cold, warmth, hard soil, loose soil, anoxia, toxins, pests, pathogens, trampling, UV radiation, fungicides, and so on – differentially changes root and shoot growth rates. It is more useful to interpret these changes not so much as specific responses to those factors, but as unavoidable consequences for the whole plant of certain of its meristems encountering them. Thinking of a plant not as an entity comprising two juxtaposed centres of growth (‘roots’ and ‘shoots’), but as populations of dispersed, multiple meristems that happen to be connected to one another, subsets of which can experience and respond separately to different local environmental conditions, removes the need for a functional equilibrium model – or any other model – of ‘adaptive’ root-shoot behaviour, at least insofar as the relationship between allocation and whole-plant growth is concerned.

It is time to move on from using the functional equilibrium model to explain what changes in root-shoot allocation mean because (a) it doesn’t really explain much and (b) it’s wrong anyway. As a metaphor to fill the gaps where experimental data should have been it stimulated us to think critically about our understanding of allocation and growth, but it has proved a dead-end for advancing that understanding any further. I offer no superior theory to replace it because none is needed.

Instead, to understand the dynamics of root-shoot biomass allocation and whole-plant growth rate it is better simply to measure root and shoot growth repeatedly and frequently (and not rely on potentially misleading ‘final harvest’ measurements), apply defined environmental treatments during growth for comparison with an untreated control so that deviations in growth or allocation between the two can be followed unambiguously (see Robinson & Peterkin 2019), and analyse temporal dynamics of allocation, growth and ideally resource capture (Trinder *et al.* 2013) using a suitable model (not necessarily the logistic but whatever is most appropriate) to help distinguish genuine responses from consequential changes in allocation and growth.

It is often true in science that ‘The basic problem…is the very common one of the easily measured variables not being the theoretically important ones’ (Williams 1966, p. 106). Here, however, the easily measured variables – root and shoot biomasses – *are* the theoretically important ones. They are measured so easily compared with many others that the potential richness of the information they contain has remained hidden and their importance overlooked. Such basic data have been analysed in too restrictive a way to realise their full explanatory value.

The production and allocation of biomass aren’t everything, of course (Kong & Fridley 2019). What really matters is what a plant does with that biomass once produced. How, and how quickly, leaves and roots take up and use raw materials, how cells metabolise assimilates, how a plant changes shape as it develops, and how it executes numerous other processes, are what a real plant does. And in a real plant feedback between roots and shoot occurs continually, not least in the upward and downward flows of vascular fluids and their contents, the activities of one plant compartment influencing those of the other. Root and shoot activities are limited also by structural and stoichiometric constraints. What effects might all these have on my conclusions? My guess is that the net effect of including them explicitly will be to restrict the occupiable RGR-RMF space in Fig. 4 to a narrower range, but not to fundamentally alter the general picture. After all, roots and shoots of real plants with all their feedback mechanisms and constraints still grow sigmoidally, on average, and that’s what the imaginary plants in Fig. 4 are doing, too.

All of that applies to understanding the dynamics of allocation and growth in individual plants as they go through their lives. What about comparisons among species? It is a biological and mathematical inevitability that no plant’s RGR is constant and highly unlikely that its RMF is either. Even so, in in multi-species screening experiments and meta-analyses RGR and RMF are averaged over many days or weeks to provide valuable but ‘static’ indices of comparative performance. Interspecific comparisons between such indices can produce puzzling or contradictory correlations between RGR and root-shoot allocation that vary in both sign and strength (Hunt & Cornelissen 1987). The explanation for this prompted by Fig. 4 is that it depends where different plants happen to be on their dynamic trajectories within the RGR-RMF landscape at the time(s) of measurement. That whereabouts is usually unknown and will always be unknown if meta-analyses don’t include temporally detailed data.

Is it true that a plant should always maximise its instantaneous growth rate anyway? That principle is often used as a goal when growth and allocation are modelled whether as isolated individuals (Thornley 1998) or as competing populations (Vincent & Vincent 1996). Most plants usually grow faster if allowed and are physiologically able. Whether that behaviour leads eventually to any fitness benefits in terms of greater seed production or vegetative spread compared with the plant’s competitors depends on many factors beyond the horizon of this paper. Any demographic advantage accruing from individuals maximising their growth rates is highly context-dependent. It seems reasonable that a faster individual growth rate relative to a competitor’s should be advantageous in the short-term, but there is no direct evidence for such an advantage. In contrast there is ample evidence for strong long-term selection against inherently fast growth in hostile habitats. There, plants typically have evolved primarily to maximise their chances of conserving resources, deterring predators and pathogens, and protecting meristems inside durable tissues, traits physiologically and developmentally incompatible with sustained fast growth (Grime 2001, p. 89).

## Concluding comments and coda

- Sigmoid growth trajectories of roots and shoots, and how these translate into dynamic changes in biomass allocation and production rates, are incompatible with a functional equilibrium model.
- Changes in allocation are caused by differential growth rates between roots and shoots. Anything that changes root or shoot growth rate will change root-shoot allocation and will lead unavoidably to a change in whole-plant growth rate.
- It will often seem as though root-shoot allocation is the primary response to the environment and so likely ‘adaptive’, but that is an illusion.
- Stronger candidates for primary, potentially ‘adaptive’ growth responses are the localised and transient productions of roots and leaves in more favourable microsites within the plant’s immediate environment. Those responses entail changes in growth rate of roots or shoots and that leads automatically to unavoidable changes in root-shoot allocation and whole-plant growth rate. The latter are secondary consequences of the primary responses.
- Biomass allocation and whole-plant growth rate are indivisible processes. They are not independent traits.
- Biomass allocation and production rate have no unique relationship to one another but can vary across a wide spectrum of possible relationships.
- Changes in root-shoot allocation cannot of themselves compensate directly for an impairment of growth rate caused by an external factor such as nitrogen shortage; such changes cannot be ‘adaptive’.

I hope by now you are thinking that the examples I used to develop my arguments are interesting as far as they go but are also asking if my focusing on just those few cases is entirely legitimate. What about all the others that I haven’t considered? Could they tell a different story? Rather than present a bestiary of all possible growth and allocation trajectories here, can I encourage you instead to open a spreadsheet or write some code to explore first-hand the sometimes surprising dynamics of biomass production and allocation via eqn. (1) along with eqns. (2) and (3)? (After writing that I remembered that May (1976) had also urged his readers to play around with eqn. (1) to best appreciate its rich counter-intuitive dynamics. It was a good suggestion then, and it is now.) Then, suitably inspired, do the experiments.

